# A bi-kinase module sensitizes and potentiates plant immune signaling

**DOI:** 10.1101/2023.03.14.532616

**Authors:** Philipp Köster, Gefeng He, Qiuyan Dong, Katarina Hake, Ina Schmitz-Thom, Paulina Heinkow, Jürgen Eirich, Lukas Wallrad, Kenji Hashimoto, Stefanie Schültke, Iris Finkemeier, Tina Romeis, Jörg Kudla

## Abstract

Systemic signaling is an essential hallmark of multicellular life. Pathogen encounter occurs locally but triggers organ-scale and organismic immune responses. In plants, elicitor perception provokes systemically expanding Ca^2+^ and H_2_O_2_ signals conferring immunity. Here, we identify a Ca^2+^ sensing bi-kinase module as becoming super-activated through mutual trans-phosphorylation and imposing synergistically enhanced NADPH oxidase activation. A combined two-layer bi-kinase/substrate phospho-code allows for sensitized signaling initiation already by near-resting elevations of Ca^2+^ concentration at the infection site. Subsequently, it facilitates further signal wave proliferation with minimal amplitude requirement, triggering protective defense responses throughout the plant. Our study reveals how plants build and perpetuate trans-cellular immune signal proliferation while avoiding disturbance of ongoing cellular signaling along the path of response dissemination.

**One sentence summary:** Mutual trans-activation of a Ca^2+^ sensing bi-kinase module potentiates NADPH oxidase activation to facilitate systemic immunity.

---

Initial to any immune response is pathogen perception by specifically committed pattern recognition receptors (PRRs) (*1*). Elicitor sensing in the primary infected cells prompts auto- and transphosphorylation of involved receptor like kinase (RLK) and receptor like protein (RLPs) complexes that subsequently trigger local and systemically spreading Ca^2+^ and *reactive oxygen species* (ROS) signals through activation of NADPH oxidases (NOX) (*2, 3*). NOX-dependent physiological generation of ROS for manifestation of systemic innate immunity is highly conserved across virtually all multicellular life (*4*). However, how distal cell-to-cell/trans-cellular propagation of these second messenger signals is perpetuated in the absence of elicitor stimulation remains largely enigmatic(*5*–*7*).

In *Arabidopsis thaliana* (hereafter Arabidopsis), perception of the bacterial elicitor peptide flg22 through the RLK FLS2 and its coreceptor BAK1 leads to phosphorylation and activation of the RLcK BIK1(*8, 9*). BIK1 functions in initiating subsequent Ca^2+^ and ROS signals (*10, 11*). BIK1 and the Ca^2+^ activated kinase CPK5 directly phosphorylate the NOX RBOHD, thereby triggering apoplastic formation of superoxide (O2-) and consequently H_2_O_2_(*12, 13*) (*14*). Accordingly, *rbohd, bik1* and *cpk5* mutants exhibit compromised local and, as consequence impaired systemic immune responses(*12*–*15*). Ca^2+^ signals are not only decoded through Ca^2+^- dependent protein kinases (CPKs), but also through a signaling network formed interaction of calcineurin B-like (CBL) Ca^2+^ sensor proteins with CBL-interacting protein kinases (CIPKs)(*16*). CBL/CIPK complexes regulate a multitude of crucial transcription factors, ion-channels and transporters and also activate the NOXs RBOHC and RBOHF (*17*–*19*). However, if and how CBL/CIPK Ca^2+^ signal sensors/decoders also function in systemic immunity signaling, remains to be addressed.

## A local Ca^2+^/phosphorylation switch triggers systemic innate immunity

We pursued a BiFC-based interaction screen, combining all 26 CIPKs from Arabidopsis with RBOHD and identified CIPK26 as most strongly interacting with RBOHD (Fig.1A). Kinase-NOX interaction occurred at the plasma membrane (PM) and active CIPK26 displayed stronger interaction than a kinase-inactive variant (Fig.1B). Recombinant CIPK26 phosphorylated the N-terminal domain of RBOHD with similar efficiency as the known CIPK26 substrate RBOHF (Fig.1C)(*17*). To address if and how CIPK26 regulates RBOHD NOX activity in a cellular context, we employed reconstitution of the plant Ca^2+^-signaling/ROS generation module in human HEK293T cells (Fig.1D). While expression of RBOHD alone only allowed for minor NOX activation, coexpression of CBL1/CIPK26 Ca^2+^ with RBOHD conferred readily detectable NOX activation already at basal Ca^2+^ concentration and was fivefold enhanced upon elicitation of Ca^2+^ signals. The most closely related kinase CIPK3 did not evoke NOX activation, indicating faithfully preserved substrate specificity in the HEK293T cell reconstitution system. RBOHD activation strictly depended on kinase activity and the presence of both, the CBL1 Ca^2+^ sensor and the kinase (Fig.1E, Fig.S1). Moreover, impairing Ca^2+^ binding of CBL1 by mutating critical EF-hands or abrogating CBL1/CIPK26 PM-targeting by mutating the CBL1 myristoylation motif attenuated ROS production back to levels observed by expression of RBOHD alone (Fig.1E, Fig.S1). Together these data identify CBL1/CIPK26 as Ca^2+^-sensor/kinase module that can bring about translation of Ca^2+^ signals into RBOHD-mediated ROS generation.

**Fig. 1.**
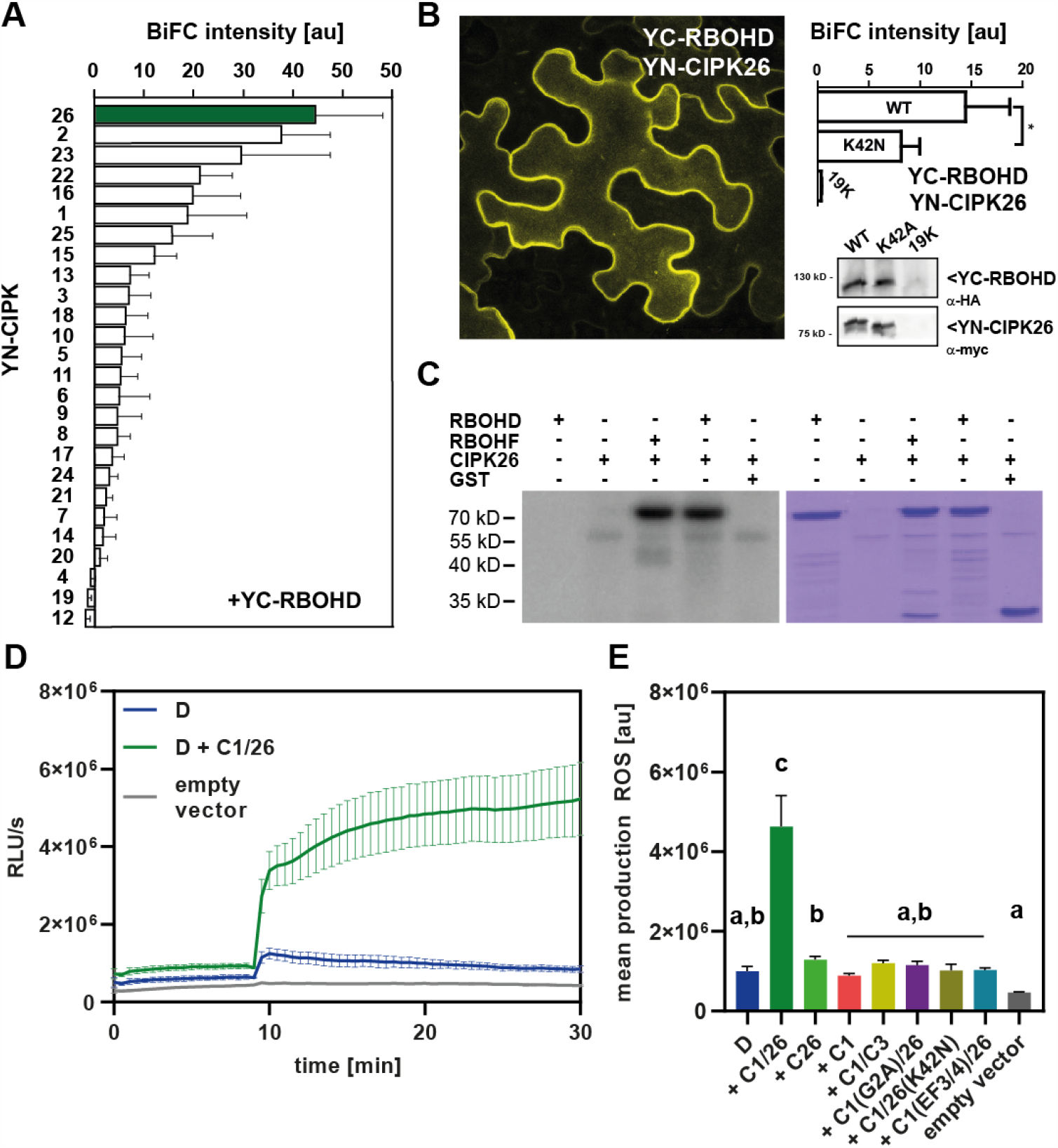
CBL1/CIPK26 associates with and activates RBOHD. (A) BiFC-based interaction screen combining each of the 26 CIPKs from Arabidopsis with RBOHD. Bars indicate mean fluorescence intensity of 6 images, error bars indicate SE. (B) BIFC analysis in *N. benthamiana* leaf epidermis cells. The image depicts a 20-stack maximum projection. Kinase-activity deficient CIPK26K42N exhibits reduced interaction with RBOHD. The bar plot illustrates mean fluorescence intensity of 6 images, error bars indicate SE. Western analysis with antibodies specific for c-myc and HA tags confirmed faithful expression of YC-RBOHD, YN-CIPK26 and YN-CIPK26K42N. (C) *In vitro* kinase assays reveal phosphorylation of the RBOHD N-terminus by CIPK26. The RBOHF N-terminus served as positive control. GST protein was not phosphorylated by CIPK26. Shown are autoradiogram and CBB stained gel. (D) CBL1/CIPK26 complexes activate RBOHD-dependent ROS production in HEK293T cells. Ca^2+^ influx into cells was initiated after 10 minutes (indicated by an arrow). ROS production was quantified as relative light units (RLU). Error bars indicate SD. Each data point represents the mean of 3 wells analyzed in parallel. (E) RBOHD activation requires CBL1/CIPK26 complex formation, CIPK26 kinase activity as well as CBL1 plasma-membrane targeting and Ca^2+^ binding (see ROS-curves in Fig. S1), one-way ANOVA, Tukey’s posttest, letters denote statistically differences between samples. **D** = RBOHD; **C1/26** = CBL1/CIPK26 complex; **C1** = CBL1; **C26(K42N)** = CIPK26K42N; **C1/3** = CBL1/CIPK3 complex; **C1(G2A)** = CBL1G2A**; C1(EF3/4)** = CBL1 EF3/4

## Collective function of CBL1/CIPK26 and CPK5 confers synergistic RBOHD activation and systemic PTI

Currently, the reasons for the coexistence of the two distinct CBL/CIPK and the CPK Ca^2+^- decoding kinase networks remain enigmatic(*16*). The identification of CPK5 and CIPK26 as kinases phosphorylating RBOHD, allowed for the first time to investigate the functional interrelation of these two kinase classes. Coexpression of CPK5 with RBOHD in HEK293T cells revealed a similar degree of Ca^2+^-dependent NOX activation as conferred by CBL1/CIPK26 (Fig.2A).

**Fig. 2.**
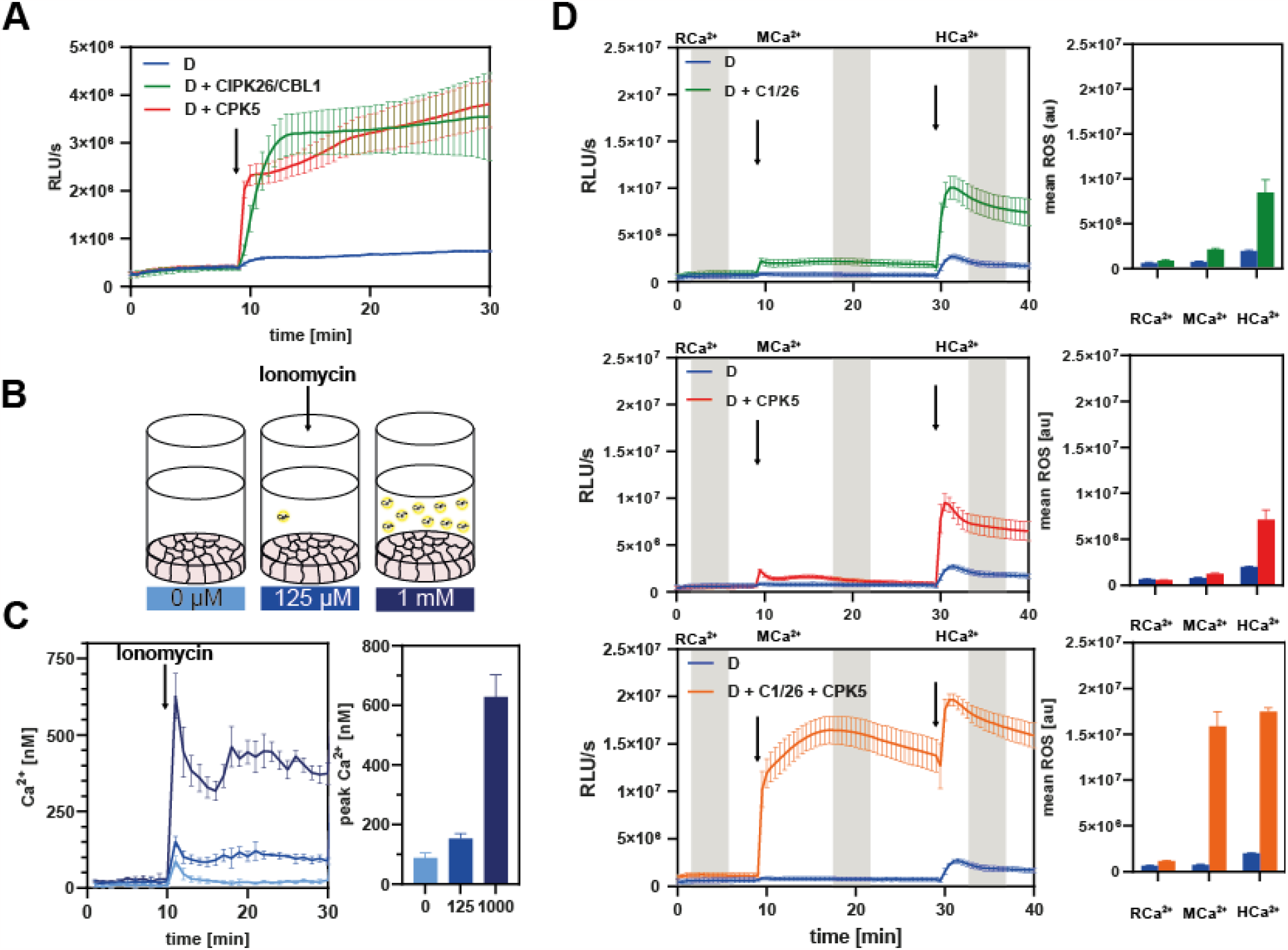
Combined activity of CPK5 and CBL1/CIPK26 triggers sensitized synergistic activation of RBOHD. (A) CPK5 and CBL1/CIPK26 similarly activate RBOHD in HEK293T cells. ROS production was quantified as relative light units (RLU). Error bars indicate SD. Each data point represents the mean of 3 wells analyzed in parallel. (B) Graphical summary of the experimental setup for controlled quantitative modulation of cytoplasmic Ca^2+^ in HEK293T cells. (C) Fura-2 based Ca^2+^ concentration determination reveals MCa^2+^ elevation to 150 nM in response to 0.125 mM external Ca^2+^ and HCa^2+^ elevation to 600 nM. Arrow indicates injection of 1 µM Ionomycin. Bar plots indicate peak Ca^2+^ elevation. Error bars indicate SD. Each data point represents the mean of 3 wells analyzed in parallel. (D) Coexpression of RBOHD with CBL1/CIPK26 and CPK5 triggers sensitized and synergistic activation of ROS production already at MCa^2+^, which is further elevated at HCa^2+^. ROS production was quantified as relative light units (RLU). Error bars indicate SD. Each data point represents the mean of 6 wells analyzed in parallel. Bar plots display mean ROS production in the 5-minute time intervals indicated by grey boxes in the depiction of the measurement curves in the left. Error bars indicate SEM.

We next sought to assess the individual roles of both kinases in RBOHD activation and to quantitatively dissect their Ca^2+^ dependence. To this end, we devised a synthetic cellular Ca^2+^- signaling reconstitution system that allowed for quantitatively controlling the amplitude of Ca^2+^ signals and thereby for faithful parameter control of regulatory circuits in HEK293T cells (Fig.2B). While ionomycin application at external Ca^2+^ concentrations of 0.125 mM triggered moderate cellular Ca^2+^ signals of approx. 150 (+/- 15) nM (moderate Ca^2+^, MCa^2+^), external Ca^2+^ concentrations of 1 mM elicited intracellular Ca^2+^ signals with an amplitude of 630 nM (+/- 60) (high Ca^2+^; HCa^2+^) (Fig.2C). Application of this protocol to cells either expressing RBOHD alone or RBOHD combined with CBL1/CIPK26 or CPK5 uncovered a strict positive correlation between cellular Ca^2+^ concentration and resulting H_2_O_2_ production indicative for RBOHD activity (Fig.2). For both CPK5 and CBL1/CIPK26, when expressed individually with RBOHD, we observed that step-wise increases in Ca^2+^ concentration incrementally increased RBOHD activity to comparable extent (Fig.2D). While MCa^2+^ triggered a 2-3x increase in H_2_O_2_ generation either with CBL1-CIPK26 or with CPK5, HCa^2+^ caused a drastically enhanced H_2_O_2_ production more than 15-fold higher compared to resting Ca^2+^ concentration (RCa^2+^, 87 nM). We next elucidated the combined effect of CBL/CIPK and CPK mediated regulation on NOX activity by co-expressing RBOHD with CBL1/CIPK26 and CPK5 simultaneously (Fig.2D, Fig.S2). HCa^2+^ triggered a notably enhanced ROS production that was doubled compared to activation by either CBL1/CIPK26 or CPK5 alone. Most remarkably, already at MCa^2+^ the combined function of both kinases caused eightfold increased NOX activity indicated by ROS production to a level that the individual kinases could not evoke even under HCa^2+^ conditions. This synergistic activation was not a consequence of increased quantity, since duplication of the amount of individually transfected kinases resulted only in additive enhancement of RBOHD activity (Fig. S2). Collectively, these findings uncover a dramatic synergistic effect of combined CPK and CBL/CIPK function on the activity of RBOHD. Moreover, these results imply simultaneous sensitization to Ca^2+^ concentration as well as potentiation of phosphorylation mediated RBOHD activation. Potential mechanisms could involve synergistic interdependences between the impact of Ca^2+^ binding to the EF-hands of CBL1, CPK5 and RBOHD (providing a combinatory Ca^2+^-code) and on the other hand, enhanced phosphorylation efficiency of p-sites either in RBOHD and/or in CBL1/CIPK26/CPK5 (providing a complementary phospho-code).

To address the physiological relevance of CBL1/CIPK26/CPK5-dependent RBOHD activation in immunity, we isolated the respective mutants and characterized their pattern triggered immune (PTI) responses and pathogen resistance. Scoring of *Pseudomonas syringae* DC 3000 proliferation three days after infiltration, indicated similarly increased bacteria abundance in *cipk26* and *cpk5* leaves and revealed substantially further enhanced bacterial proliferation in leaves of *cipk26/cpk5* (Fig.3A). Flg22 application evoked only strongly diminished local ROS accumulation in directly flg22 exposed leaf discs of *cpk5* as well as of two independent *cipk26* alleles (Fig.3B + Fig.S3). This local immune response was not further reduced in *cipk26/cpk5*, but fully abolished in *rbohD*. Flg22 application triggers induction of the marker gene *NHL10* locally, but also in distal leaves that have not experienced direct elicitor exposure(*14*). We found that local *NHL10* upregulation in flg22 challenged leaves was not impaired in *cipk26* but significantly reduced in *cipk26/cpk5* (Fig.3C). Moreover, these analyses confirmed the reported contribution of CPK5 to local NHL10 induction(*14*). In stark difference, loss of CIPK26 notably reduced *NHL10* upregulation in distal leaves, and in *cipk26/cpk5* this response was almost abolished. Collectively, these *in planta* investigations reveal an important individual role of CPK5 and CIPK26 in pathogen resistance and establish the indispensable requirement of their combined function for systemic immune signaling.

**Fig. 3.**
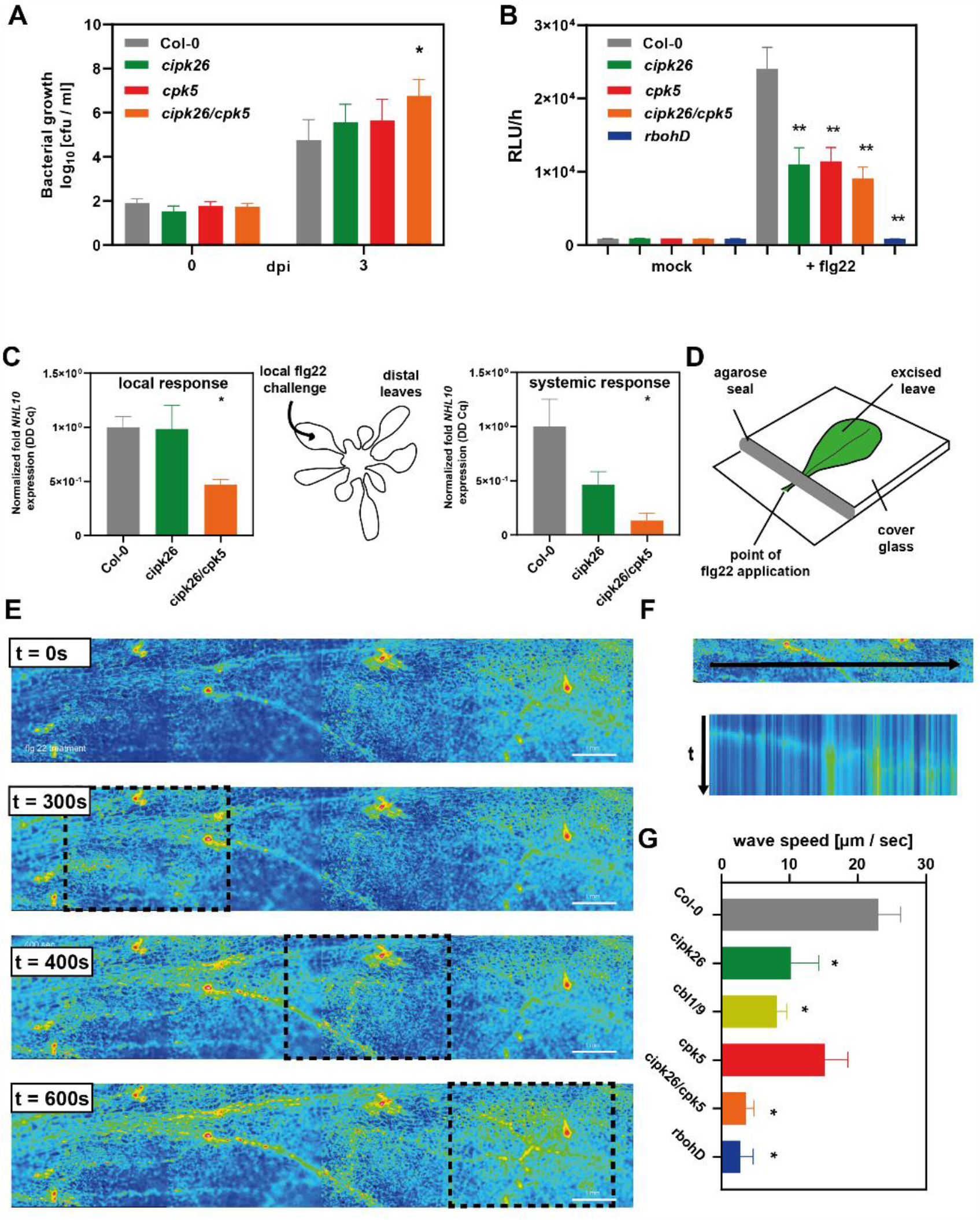
The CBL/CIPK26/CPK5/RBOHD module confers initiation and propagation of systemic immune signaling. (A) Loss of CIPK26 and/or CPK5 function renders Arabidopsis plants susceptible to *Pseudomonas syringae*. Bacterial growth was assayed in 6-week-old Col-0, *cipk26, cpk5*, and *cipk26/cpk5* plants at day 0 and 3 days post inoculation with *Pst* DC3000. c.f.u., colony forming units; error bars, SEM (n ≥ 9); one-way ANOVA, Tukey posttest, asterisks denote statistically differences compared to Col-0, * P < 0.05, **P < 0.01, **P < 0.001) (B) CIPK26 and CPK5 are required for flg22 induced respiratory burst. ROS-production was determined over 60 min via a luminol-based assay with and without treatment with 200 nM flg22 in 6-week-old plants of Col-0, *cipk26, cpk5, cipk26/cpk5*, and *rbohd*. RLU, relative light units; error bars, SEM (n ≥ 8); one-way ANOVA, Dunnett posttest, asterisks denote statistically differences compared to Col-0, **P < 0.01). (C) Flg22-induced NHL10 expression induction in distal leaves depends on CIPK26 function. Forty-five minutes after 200 nM flg22 injection NHL10 gene expression was quantified by qRT-PCR in an upper leaf. Error bars, SEM (n ≥ 7); Student t test,*P < 0.05. (D) Schematic depiction of the experimental setup for organ-scale Ca^2+^ dynamics analysis in whole leaves in response to flg22 petiole application. Leaves of Arabidopsis plants expressing the Ca^2+^ reporter proteins were detached, 24 h later flg22 was added to the petioles. The formation and dispersion of Ca^2+^ signals in the leaf blade was determined by fluorescence microscopy and the speed of the signals was measured. (E) Images of R-GECO fluorescence in an Arabidopsis leaf blade at different timepoints after flg22 application to the petiole base. Boxes highlight the progressing Ca^2+^ wave. (F) Kymograph displaying the propagation of the wave front through the indicated area line. (G) Speed of the organ-scale Ca^2+^ waves in indicated genotypes. n > 5, error bars indicate SEM. one-way ANOVA, Tukey’s posttest, asterisks denote statistically differences compared to Col-0.

We next sought to use the flg22-triggered systemic Ca^2+^ wave as a proxy to illuminate the role of Ca^2+^/phosphorylation-mediated NOX activation in systemic signaling. To this end, we devised a systemic bioimaging assay for whole detached leaves based on flg22 application onto petioles and Ca^2+^ reporter-based monitoring of the resulting Ca^2+^ waves throughout the leaf blade (Fig.3D). In WT, application of flg22 induced a Ca^2+^ wave initiating from the leaf basis, subsequently expanding to the whole width of the leaf, and forming a front that travelled towards the leaf tip (Fig.3E, F+ MovieS1). The amplitude (signal intensity) of this wave was reproducibly clearly less pronounced than that of recently described Ca^2+^ waves e.g. in response to NaCl stress(*20*). We determined a speed of 21 +/- 5 [µM/s] for systemic Ca^2+^ signal propagation through non-vascular epidermal cells in WT (Fig.3F). The speed of this newly identified flg22 response is significantly slower compared to 200-1000 µM/s that have been determined for systemic electrical, ROS or Ca^2+^ signals propagating through vascular bundles in response to other stimuli(*20*–*27*). Therefore, this value likely defines the cell-to-cell transcellular propagation velocity of PAMP triggered Ca^2+^/ROS signals. In *rbohD* the amplitude of the propagating Ca^2+^ signal was dramatically reduced and wave propagation ceased almost completely, corroborating the essential role of this NOX in cell-to-cell communication (Fig.3G). In individual mutants of CPK5, CIPK26 or CBL1/9 (the two Ca^2+^ sensors that activate CIPK26) both the amplitude and the speed of the mobile Ca^2+^ signal was significantly reduced. Notably, impairment of CIPK-mediated phosphorylation exerted a stronger effect than compromised CPK phosphorylation. Strikingly, in *cpk5*/*cipk26* systemic Ca^2+^ signal propagation was similarly collapsed as in *rbohD*. This uncovers an absolute requirement of Ca^2+^-dependent NOX phosphorylation for systemic Ca^2+^ signal formation and propagation. Moreover, since CPK and CBL/CIPK activity towards RBOHD are strictly Ca^2+^ dependent, we also conclude that this mobile Ca^2+^ signal is essential for forming the PAMP-triggered ROS wave and for evoking distal transcriptional responses like *NHL10* induction.

## Trans-acting mutual CPK5-CIPK26 phosphorylation together with cis-acting phosphorylation in RBOHD convey a two-layer activation mechanism

To dissect the mechanistic principles underlying the sensitized and potentiated activation of RBOHD, we combined control of cytoplasmic Ca^2+^ concentration with mass-spectrometric phosphorylation-site identification and quantitation. HEK293T cells expressing RBOHD alone, RBOHD combined with individual CPK5 or CBL1/CIPK26 or BIK1, or RBOHD together with CBL1/CIPK26 and CPK5 were ionomycin treated to evoke MCa^2+^ cytoplasmic Ca^2+^ signals (150 nM) to cover the conditions of sensitized hyperactivation of RBOHD. Then total protein was extracted and subjected to phospho-proteomic analysis. This approach identified seven serines and one threonine in RBOHD located on 6 distinct peptides (hereafter “p-sites”) as being principally phosphorylated upon kinase exposure (Fig. 4A, Table S1). One p-site was almost exclusively targeted by BIK1 (Ser 39), while the other p-sites were targeted by all three kinases to varying degree. Quantitative evaluation of the phosphorylation profile defined three distinct target site groups: (i) preferentially BIK1-addressed p-sites (Ca^2+^-independent: Ser 39; Ser 339), (ii) preferentially CBL1/CIPK26/CPK5-addressed p-sites (Ca^2+^-dependent: Ser 8; Ser162/163; Thr699) and (iii) p-sites concurrently targeted by BIK1 as well as CBL1/CIPK26/CPK5 (Ser 343/347). To address the functional significance of Ca^2+^-dependent phosphorylation, we elucidated the activatability of RBOHD^Ser8/162/163/Thr699A^ by CBL1/CIPK26/ CPK5. Notably, in both MCa^2+^ and HCa^2+^ conditions, H_2_O_2_ production conferred by RBOHD^Ser8/162/163/Thr699A^ was dramatically reduced to less than half of that of RBOHD (Fig. 4B). In contrast, this mutant version of RBOHD was not impaired in BIK1 mediated activation thereby establishing the independent relevance of Ca^2+^-dependent phosphorylation for RBOHD activity modulation (Fig.4B). Somewhat unexpectedly, mutation of BIK1-specific residues (RBOHD^S39,339A^) did not impact on RBOHD activatability by BIK1 or CBL1/CIPK26/CPK5 revealing that modification (or modification-preventing mutation) of these p-sites is dispensable for RBOHD activity modulation (Fig.4C). Interestingly, we discovered a surprising complex ROS generation pattern when analyzing the impact of the shared p-sites (RBOHD^S343/347A^). When RBOHD^S343/347A^ was combined with CBL1/CIPK26/CPK5, ROS production at HCa^2+^ (630 nM) was reduced by about 40% to a level produced by non-mutated RBOHD at MCa^2+^ (150 nM). Even more strikingly, at MCa^2+^ RBOHD^S343/347A^ produced only as much H_2_O_2_ as RBOHD alone (without any kinase) would produce at HCa^2+^ (Fig.4D). This identifies Ser343/347 as mechanistic switch essential for allowing sensitized activation by Ca^2+^-dependent kinases at moderate Ca^2+^ signal intensity. Also, remarkably, non-phosphorylatability of Ser343/347 rendered activation of RBOHD by BIK1 Ca^2+^ sensitive (Fig.4D). While RBOHD^S343/347A^ activation by BIK1 was already significantly reduced at HCa^2+^, this activation level collapsed down to approx. 20% of that of RBOHD at MCa^2+^ and was almost abolished at RCa^2+^. These findings reveal that Ser343/347 functions as switch that, when phosphorylated allows for maximal activation of RBOHD at resting Ca^2+^ concentration (RCa^2+^, 87 nM) solely through phosphorylation of other p-sites, while non-phosphorylated Ser343/347 necessitates higher Ca^2+^ concentrations (HCa^2+^, 630 nM) for efficient Ca^2+^ binding to RBOHD EF-hands as alternative means allowing for full NOX activation. In conclusion, the phosphorylation status of Ser343/347 likely regulates the Ca^2+^ binding affinity/efficiency of the adjacent EF-hands. Phosphorylation of Ser343/347 by either BIK1 or CBL1/CIPK26/CPK5 would therefore be a key cis-regulatory step on the substrate level for enabling Ca^2+^-sensitized ROS generation. Astonishingly, we also discovered that Ser 343/347 as well as all other p-sites that are shared between CBL1/CIPK26 and CPK5 displayed significantly enhanced phosphorylation intensity when simultaneously exposed to both kinases as compared to the phosphorylation intensity conferred by individual kinases (Fig.4A).

**Fig. 4.**
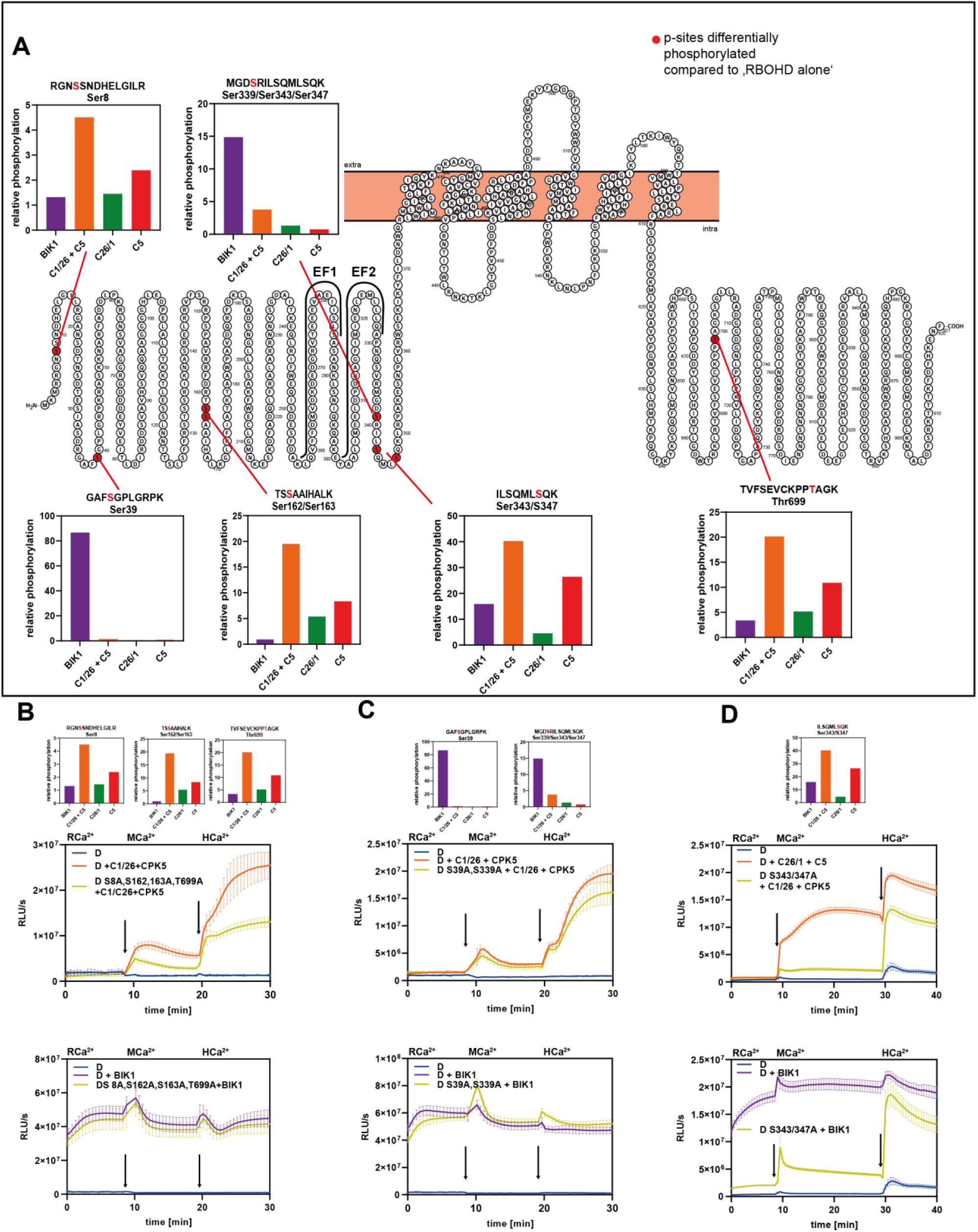
Ca^2+^-independent phosphorylation through BIK1 and Ca^2+^-dependent phosphorylation through CBL1/CIPK26 and CPK5 provide alternative routes for RBOHD activation. (A) Quantitative presentation and schematic depiction of p-sites identified as being differentially phosphorylated in response to moderate Ca^2+^ signals (MCa^2+^) in HEK293T cells expressing the indicated kinase combinations (compared to cells expressing RBOHD alone). (B) Mutation of the p-sites targeted by the Ca^2+^-dependent kinases (Ser 8, Ser162/163, Thr699) impairs activation of RBOHD by CPIK26/CPK5 but not by BIK1 in HEK293T cells ROS production was quantified as relative light units (RLU). Error bars indicate SD. Each data point represents the mean of 5 wells analyzed in parallel. (C) Mutation of the p-sites targeted by BIK1 (Ser39 and Ser339) does not impair activation of RBOHD in HEK293T cells. ROS production was quantified as relative light units (RLU). Error bars indicate SD. Each data point represents the mean of 5 wells analyzed in parallel. (D) Mutation of the p-sites shared by CIPK26/CPK5 and BIK1 (Ser343/347) render BIK1-mediated RBOHD activation Ca^2+^-dependent and abolish Ca^2+^- phosphorylation-dependent activation of RBOHD. ROS production was quantified as relative light units (RLU). Error bars indicate SD. Each data point represents the mean of 6 wells analyzed in parallel.

We therefore considered, that CBL1/CIPK26 and CPK5 could mutually enhance their capability for substrate phosphorylation by e.g., reciprocal trans-phosphorylation. Indeed, inspection of our phospho-proteomic data set identified Ser158/161 in CIPK26 as being principally phosphorylated and displaying a higher degree of phosphorylation upon co-expression with CPK5 (Fig.5A). Also, in CPK5, we found that intensity of Ser337/338 phosphorylation was enhanced by CBL1/CIPK26 coexpression (Fig.5A). Indeed, *in vitro* phosphorylation assays combining inactive CIPK26 (CIPK26^K42N^) with active CPK5 established that CPK5 efficiently trans-phosphorylated CIPK26 (Fig.5B). Vice versa, phosphorylation of inactive CPK5 (CPK5^D221A^) by active CIPK26 was also readily detectable (Fig.5B). To molecularly detail the contribution of Ser158/161 (in CIPK26) and Ser337/338 (in CPK5) to conferring Ca^2+^ sensitization of RBOHD activation, we characterized their potential to activate this NOX at MCa^2+^. Mutation of Ser158/162 in CIPK26 (CIPK26^S158/161A^) did not significantly impact on its ability of CPK5 to activate RBOHD, while mutation of Ser337/338 (CPK5^S337/338A^) reduced the individual capability of CPK5 to activate RBOHD (Fig.5C). However, when we assayed different combinations of mutated and WT kinases in HEK293T cells, we discovered trans-phosphorylation of both kinases as being required for Ca^2+^-sensitized RBOHD activation (Fig.5D). Reciprocal combination of non-phosphorylatable with phosphorylatable kinases (CPK5^S337/338A^ + CBL1/CIPK26 and CPK5 + CBL1/CIPK26^S158/161A^) both dramatically reduced the synergistic activation of RBOHD when compared to combination of mutually trans-phosphorylatable kinases (CPK5 + CBL1/CIPK26). Here, the effect of CPK5^S337/338^ phosphorylatability appeared to be more pronounced, coinciding with the high Ca^2+^ responsiveness (KD_50_ 100 nM) of CPK5 (*15*). Remarkably, combination of both mutant versions (CPK5^S337/338A^ + CBL1/CIPK26^S158/161A^) even further diminished the synergistic RBOHD activation capability normally conferred by collective function of both kinases.

**Fig. 5.**
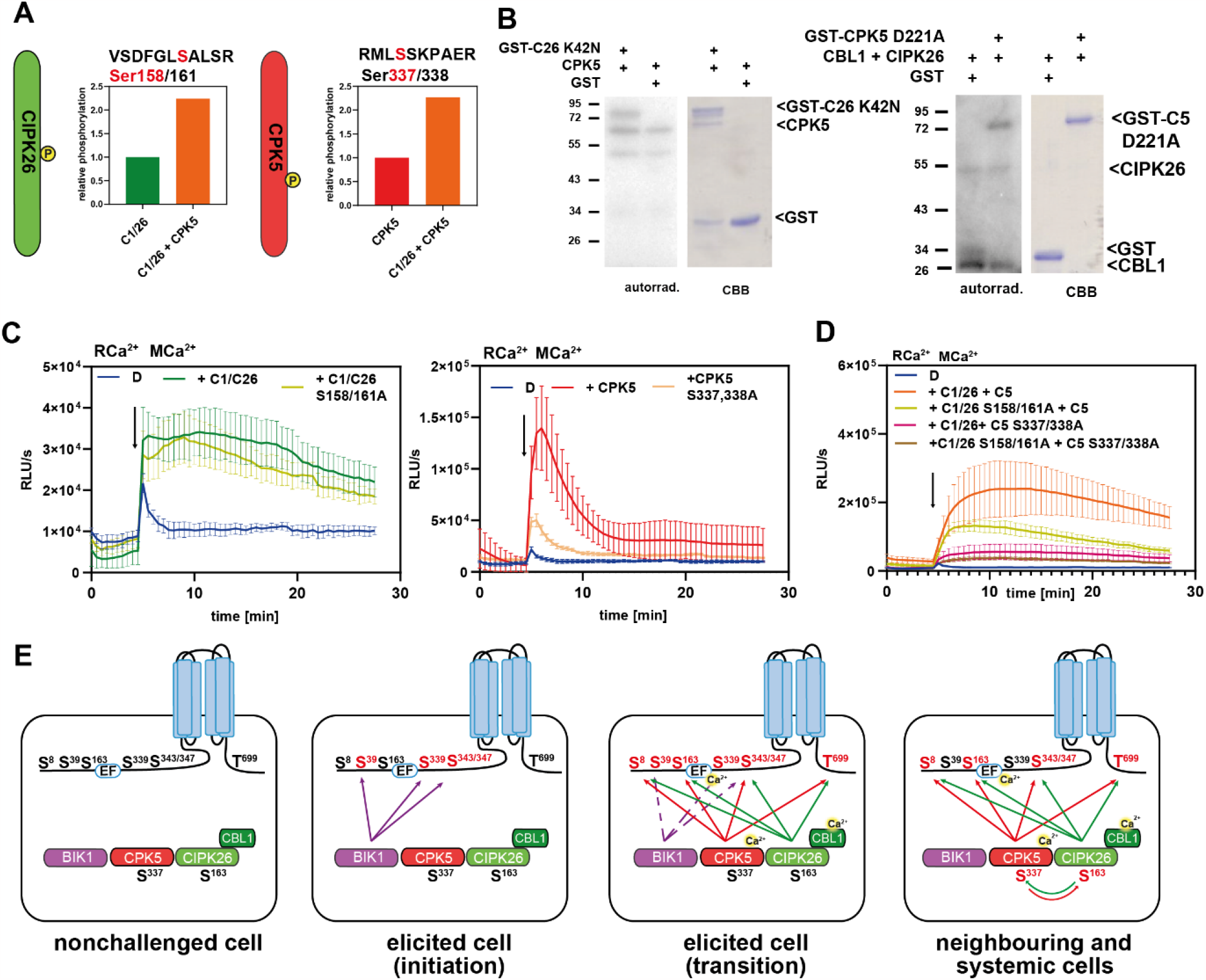
Mutual hyperactivation of CIPK26 and CPK5 through trans-phosphorylation and integrated cellular- and molecular-scale model for Ca^2+^/ROS-mediated systemic immune signaling. (A) S in CIPK26 and S in CPK5 represent differentially transphosphorylation p-sites. The bar graphs indicate phosphorylation intensity determined in HEK293T cells expressing the combination of both kinases compared to cells expressing the individual kinases at MCa^2+^. (B) CPK5 phosphorylates kinase-inactive CIPK26 *in vitro* and *vice versa*. Depicted are autoradiograms and CBB stained gels. (C) Impact of p-site mutation on the individual ability of CIPK26 and CPK5 to activate RBOHD. ROS production was quantified as relative light units (RLU). Error bars indicate SD. Each data point represents the mean of 6 wells analyzed in parallel. (D) Impact of p-site mutation on the combined ability of CIPK26 and CPK5 to activate RBOHD. ROS production was quantified as relative light units (RLU). Error bars indicate SD. Each data point represents the mean of 6 wells analyzed in parallel. (E) Molecular model of the phosphorylation events forming the combined phospho- and Ca^2+^-codes underlying systemic immunity.

## Emerging principles of Ca^2+^ sensitized RBOHD activation in systemic PTI

Systemic signaling in multicellular organisms involves organ-scale (trans-cellular) as well as organismic (trans-organ) proliferation mechanisms. In plants, organismic signals transduce via vasculature wires and mechanistically involve electrical, chemical and mechanical components, while in animals nerve cords and vasculature provide convergent solutions to serve the dissemination of electrical or hormonal signals(*28*). Trans-cellular signaling within an organ involves diffusion and/or propagation of second messengers and can serve dispersion of incoming signals within an organ but can also precede subsequent trans-organ signal dissemination(*24, 27*). Mutually interdependent Ca^2+^ and ROS waves crucially function in both organismic and organ-scale signaling(*7, 21, 23*). Here we identify and characterize a tri-molecular CBL1/CIPK26/CPK5 module as functioning in both processes through simultaneously conferring Ca^2+^-sensing and Ca^2+^-dependent phosphorylation for activating the NADPH oxidase RBOHD, a key enzyme for mounting plant immunity and stress tolerance. Combining two distinct kinase families in one dual-Ca^2+^-sensing and dual-phosphorylating module tremendously expands signaling versatility by extending the range of addressable target motifs and enlarging the concentration range of decodable Ca^2+^ signals.

A first important insight emerging from our studies, is that all components of this tri-molecular signaling module crucially contribute to organismic implementation of systemic immunity, because representative PTI responses are impaired in distal leaves after local elicitor application in individual *cipk26* and *cpk5* mutants. Abolishment of PTI in *cipk26/cpk5* establishes the absolute requirement of this module for interorgan proliferation of systemic immunity. The combined outcome of our marker gene induction analysis and Ca^2+^ signaling assays specify the binary CIPK26/CPK5 kinase module as conferring both, local (trans-cellular) and systemic (trans-organ) immune responses. We further focused on detailing the molecular mechanisms that define the function of this tri-molecular signaling module in organ-scale manifestation of trans-cellular Ca^2+^/ROS signal propagation. A striking insight with profound conceptual consequences emanating from these investigations is that the PRR-activated but Ca^2+^-independent kinase BIK1 and the Ca^2+^-activated kinases CBL1/CIPK26 and CPK5 confer phosphorylation of distinct (“Ca^2+^-independent” and “Ca^2+^-dependent”) but also of a shared set of target sites. Of these, the shared Ser343/347 p-site apparently brings about cis-acting Ca^2+^ sensitization of RBOHD. Our experimental data do not allow to distinguish if this results from modulation of the Ca^2+^ binding efficiency of the EF-hands in RBOHD or if the Ser343/347 phosphorylation status brings about a conformational change mimicking Ca^2+^ binding to these EF-hands. In line with a central role of Ser343/347 as switch for sensitizing RBOHD activation is that Ser 343/347 are the most strongly phosphorylated residues in the whole Arabidopsis proteome after flg22 treatment(*29*). Another most relevant discovery reported here is the mutual trans-phosphorylation of CPK5 and CIPK26, which causes synergistically enhanced RBOHD substrate activation of both kinases already in response to delicate Ca^2+^ signals (MCa^2+^; 150 nM). This trans-acting Ca^2+^ sensitization mechanism occurs at concentrations close to the resting level of the cytoplasm (RCa^2+^; 87 nM) and coincides with the high Ca^2+^ sensitivity of CPK5 (KD_50_ 100 nM). The critical Ser337/338 site in CPK5 resides in a loop adjacent to the autoinhibitory pseudosubstrate-domain of this kinase, while in CIPK26 the critical Ser158 residue locates in the DFG +2 position of the activation loop. Remarkably, previous studies on mammalian and yeast kinases have established that the identity and phosphorylation of the respective residues in these kinases modulates their substrate phosphorylation and preference, suggesting a potential mechanism for the trans-acting mutual activation of CPK5 and CIPK26(*30, 31*). This discovery solves the long-standing enigma, of how trans-cellular second messenger waves can propagate through organs without interfering with intra-cellular signaling processes in individual cells along this path. The combined consequence of this two-layer Ca^2+^ sensitization and activity potentiation mechanism reported here allows for organ-scale dispersion of a low amplitude Ca^2+^ wave (that we detected here in flg22 treated leaves) by “sidling” through individual cells without disturbing ongoing essential intracellular Ca^2+^ signaling processes. Notably, the trans-activating p-sites in CIPK26 and CPK5 are not unique to these kinases, but appear to be conserved in several, but not all, members of both kinase families (Fig. S4). The apparent conservation of this trans-sensitization/activation mechanism is suggestive for the occurrence of similar mechanisms in other systemic stress response processes in plants.

Collectively, our findings allow to deduce a model for switching from initial local, to subsequent systemic signaling in innate immunity, and for sustaining this long-distance signal after initiation (Fig.5E). In the primary elicitor exposed cell(s), elicitor binding activates the PRR complex and confers direct activation of specific RLcKs, including BIK1(*12, 13*). During defense signaling initiation, BIK1 in turn phosphorylates RBOHD at multiple sites including Ser343/347 resulting in NOX activation and local extracellular ROS production. These directly elicitor-stimulated cells also form primary Ca^2+^ signals that may or may not contribute to RBOHD activity via direct EF-hand binding and activation of Ca^2+^ dependent kinases. Paracrine signaling by apoplastic ROS activates Ca^2+^ channels in neighboring cells, allowing for subtle increases in cellular Ca^2+^ concentration, that suffice to activate the CBL1/CIPK26/CPK5 module in the absence of PRR activation. Both kinases synergistically phosphorylate and thereby activate RBOHD for maximal ROS production already triggered through minute elevation of cytoplasmic Ca^2+^ concentration. This allows signal propagation to the next distal cell eventually forming an iterative paracrine cell-to-cell signaling circuit manifesting as propagating Ca^2+^/ROS signal. In this way, the CBL1/CIPK26/CPK5/RBOHD module concomitantly confers organ-scale signaling initially within the primary challenged leaf, but also subsequently throughout the whole plant resulting in systemic immunity.

## Acknowledgements

We thank A. von Schaewen for providing access to the Tecan Sapphire 2 plate reader. This research was supported by the Deutsche Forschungsgemeinschaft (DFG) (Research Unit FOR964) to J.K. and T.R. and the Collaborative Research Centre SFB973 to T.R as well as the grants 290476080 and 469950637 to IF.

## Author Contributions

J.K. conceived, and J.K. and T.R. supervised the research. P.K., G.H., Q.D., K.H., I.S.-T., P.H., J.E., L.W., K.H. and S.S. performed experiments and together with I.F., T.R. and J.K. analyzed the data. P.K. and G.H. prepared the figures. J.K. and P.K. wrote the manuscript and all authors contributed.

---

**Fig. S1.**
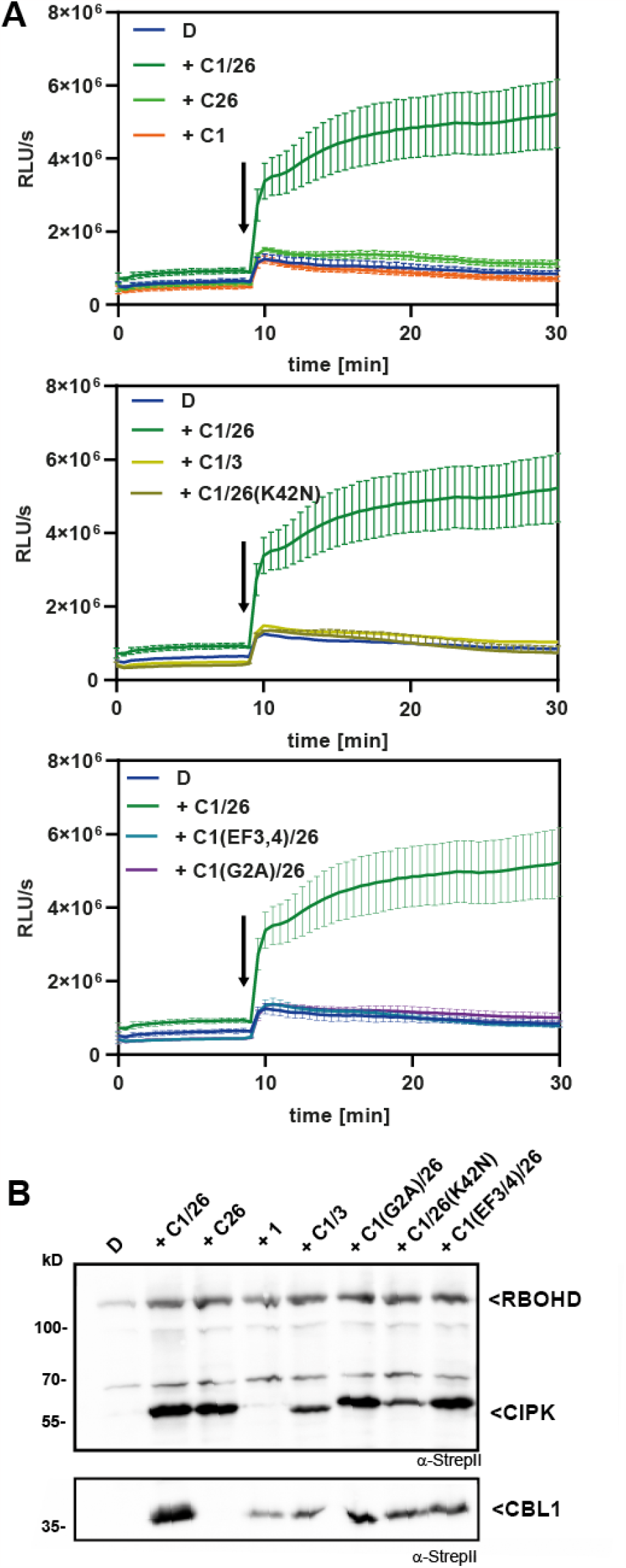
Kinase activity of CIPK26, CBL1 membrane targeting and Ca^2+^-binding are required for RBOHD activation. Displayed are individual curves of the samples shown in Fig.1D. Ca^2+^ influx into cells was initiated after 10 minutes (indicated by an arrow). ROS production was quantified as relative light units (RLU). Error bars indicate SD. Each data point represents the mean of 3 wells analyzed in parallel. (B) Western blot indicating expression of the respective proteins from the HEK293T cells assayed in Fig.1D. Total protein extract was analyzed by western blot, StrepII-tagged proteins were visualized using α-StrepII antibody. **D** = RBOHD; **C1/26** = CBL1/CIPK26 complex; **C1** = CBL1; **C26(K42N)** = CIPK26K42N; **C1/3** = CBL1/CIPK3 complex; **C1(G2A)** = CBL1G2A**; C1(EF3/4)** = CBL1 EF3/4

**Fig. S2.**
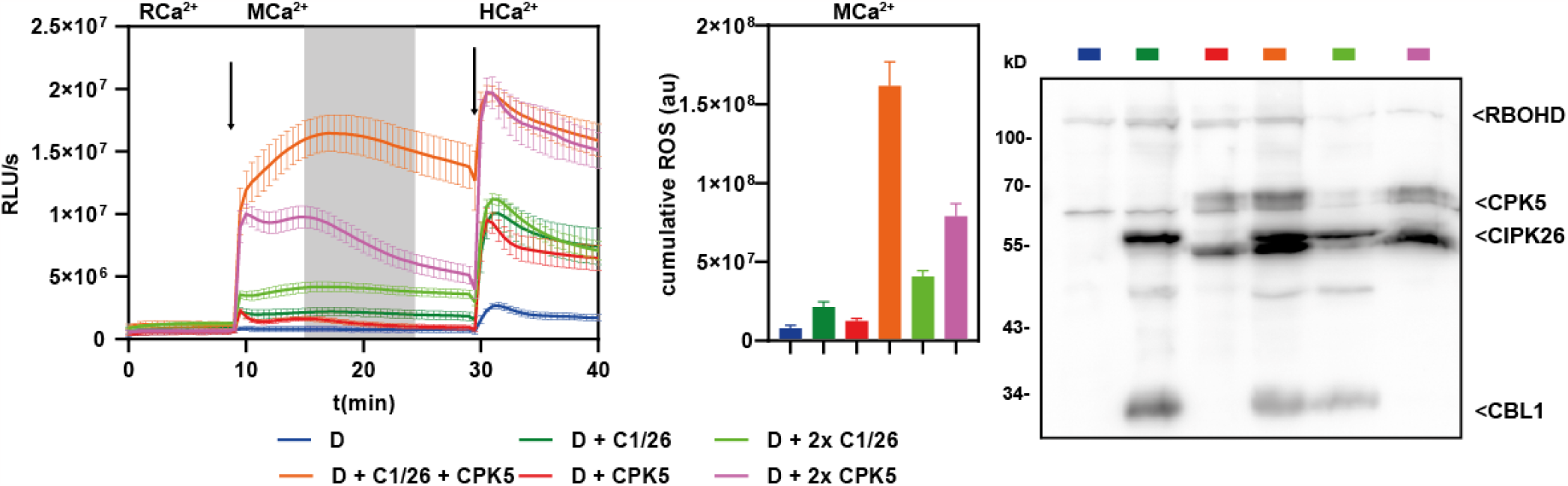
Synergistic activation of RBOHD by CBL1/CIPK26 and CPK5 does not result from increased expression of individual kinases. Doubling the amount of transfected plasmid DNA of either CBL1/CIPK26 or CPK5 results in additive but not synergistic activation of RBOHD. The measurement curves for D, D + C1/26, D + CPK5, D + C1/26 + CPK5 are identical to Fig.2D since they result from the same experiment. Western blot indicating expression of the respective proteins from the HEK293T cells assayed in Fig.2D. Total protein extract was analyzed by western blot, StrepII-tagged proteins were visualized using α-StrepII antibody. **D** = RBOHD; **C1/26** = CBL1/CIPK26 complexes.

**Fig. S3.**
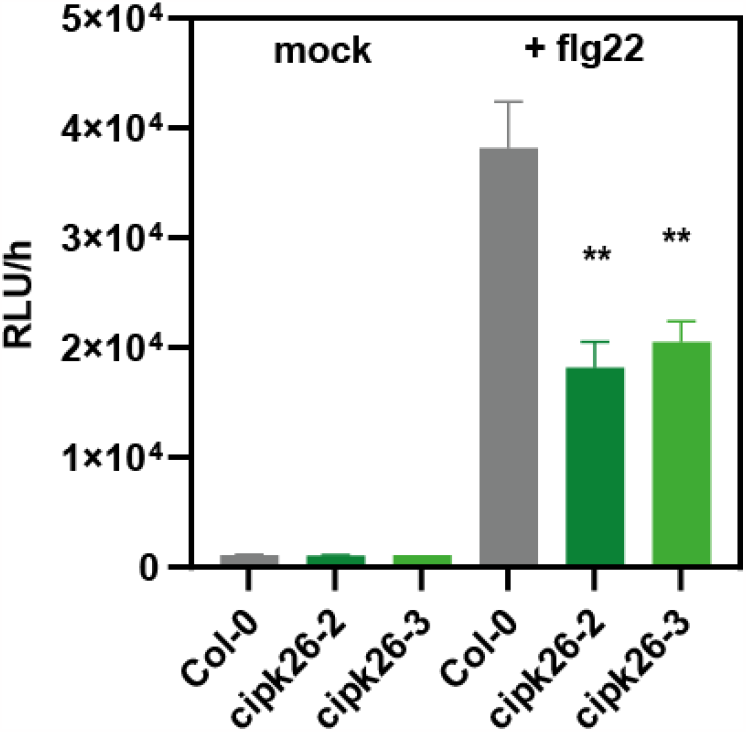
Two independent mutant alleles of *CIPK26* are impaired in flg22 induced ROS generation. ROS generation was determined over 60 min via a luminol-based assay with and without treatment with 200 nM flg22 in 6-week-old plants of Col-0, *cipk26-2*, and *cipk26-3*. RLU, relative light units; error bars, SEM (n = 16); one-way ANOVA, Dunnett posttest, asterisks denote statistically differences compared to Col-0, **P < 0.01).

**Figure S4:**
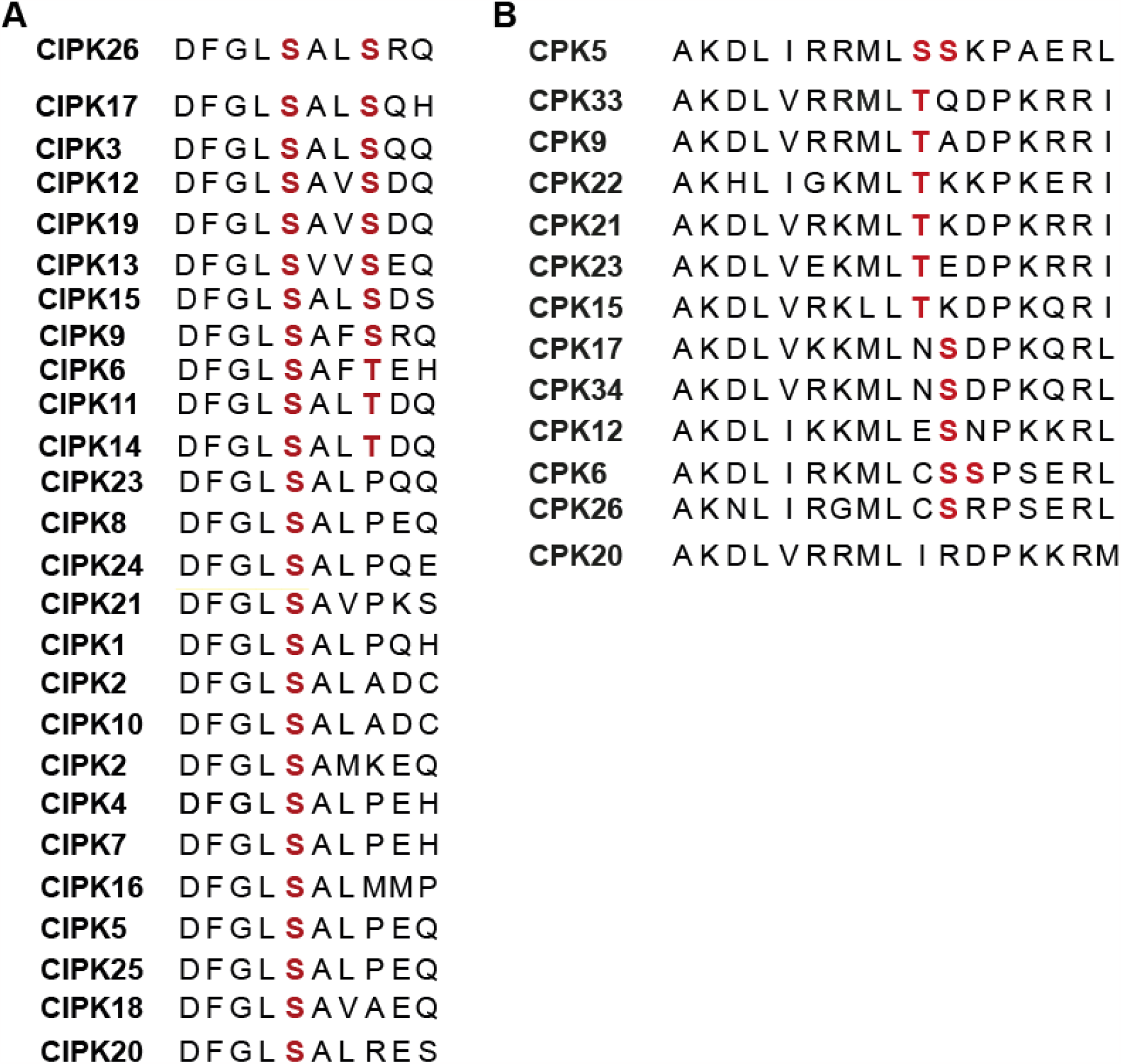
Alignments of the activating p-sites detected in CIPK26 and CPK5. p-sites in the respective regions within CIPK26 and CPK5 are highlighted in red. CIPK26 Ser158 is strictly conserved in Arabidopsis CIPKs (A), in Arabidopsis CPKs Ser and Thr residues at the respective positions of CPK5 Ser337 and Ser338 are conserved in 12 of 34 isoforms.

**Table S1: Phospho-peptides found in CBL1, CIPK26, CPK5, BIK1 and RBOHD proteins after heterologous expression in HEK293T cells and enrichment.** For each sample, the intensities detected in each of the four independent replicates are provided.

## Material and Methods

### Plant materials and growth conditions

*Arabidopsis thaliana* ecotype Columbia-0 (Col-0) was used as wild type and T-DNA insertion lines *cipk26*-2 (GK-703D04) and *cipk26*-3 (SALK_005859C), *rbohd* (SALK_070610) and *cpk5* (SAIL_657_C06) were obtained from the European Arabidopsis Stock Centre (http://arabidopsis.info). *cipk26-2* (GK-703D04) and *cipk26-3* as well as *cbl1/9* were described earlier(*1*–*3*). *Arabidopsis thaliana* Col-0 wild-type and derived transgenic overexpressing and mutant plants were grown on soil with 8 h light/16 h dark cycle, 23 °C and 60 % relative humidity. *Nicotiana benthamiana* plants were grown on soil in a greenhouse with 16 h light/8 h dark cycle.

### Bacterial growth assays

Bacterial pathogen *Pseudomonas syringae* pv. tomato DC3000 was grown in King’s B medium at 28 °C overnight. For measuring bacterial growth, Arabidopsis leaves of six-week old plants were infiltrated with Pst DC3000 at 104 cfu/ml in 10 mM MgCl_2_ using a needleless syringe. Three days after inoculation, bacterial growth was monitored by serial dilution plating of ground leaf discs.

### ROS measurements in Arabidopsis

Reactive oxygen species production was monitored using a luminol-based assay. Flagellin-dependent oxidative burst in *A. thaliana* was conducted with six-week old plants grown under short day conditions. 0.3 cm leaf discs were floated overnight on 100 µl H_2_O in a 96-well plate.

### Gene expression by qRT-PCR analysis

To analyze transcript levels, RNA was extracted from leaf tissue using the Trizol method. 2 µg of RNA was treated with RNase-free DNase (Fermentas) and reverse transcribed with SuperscriptIII SuperMix (Invitrogen) according to the manufacturer’s protocols. Real-time quantitative PCR analysis was performed in a final volume of 10 µl according to the instructions of Power SYBR Green PCR Master Mix (Applied Biosystems) using the CFX96 system (Bio-Rad). Amplification specificity was evaluated by post-amplification dissociation curves. ACTIN2 (At3g18780) was used as the internal control for quantification of gene expression.

### Calcium imaging

#### Growth conditions and sample preparation

For the analysis of Ca^2+^ signals evoked by flg22 in leaves, Arabidopsis plants were germinated on MS solid media and transferred to soil approx. 1 week after germination and grown under long-day conditions. True leaves of approx. 1 cm length were cut off and incubated for 24 hours under constant light in incubation buffer (10 mM MES, 5 mM KCl, 10 mM CaCl_2_, pH adjusted to 5.8 with Tris). Subsequently, leaves were mounted upside-down on a microscope slide. At the petiole, a twofold barrier consisting of 1% low melting point agarose and a second layer of plasticine was created to separate the leaf blade from the petiole (see Fig.4B). The purpose of this barrier is to prevent access of the elicitor later added to the petiole. Finally, the sample was covered with a cover-slide. After this mounting procedure, samples were again incubated for at least 6 hours under continuous light.

#### Epi-fluorescence image acquisition for measurement of calcium dynamics

For R-GECO1-mTurquoise-based in vivo Ca^2+^ imaging(*4*), an inverted ZEISS Axio observer microscope was used (Carl Zeiss Microimaging GmbH, Goettingen, Germany), which was equipped with a xenon short arc reflector lamp (Hamamatsu) a Zeiss EC Plan-NEOFLUAR 5x/0.16 dry objective, an ET436-20x T455lp ET480-40m filter set for mTurquosise, an ET560-40x T585lpxr ET630-75m filter set for R-GECO1, and a Retiga R6 camera, and operated by the Visiview software (Visitron Systems GmbH, Puchheim Germany).

An exposure time of 400 ms was used for both R-GECO1 and mTurquosise image acquisition with binning 2. Four areas of the leaves were consecutively analyzed every 10 seconds to cover the majority of the leaf area. After measurements for 20 cycles of 10 seconds each, 10 µL of a 4 µM flg22 solution was pipetted to the petiole. Ca^2+^ measurements were then proceeded for 30 minutes. For each measurement, ratio images of the four leaf areas were combined to ratio stacks and finally the four ratio stacks were combined to a single stack using ImageJ. To determine the speed of the Ca^2+^ waves, ROIs with approximate size of an epidermal cell were defined in the proximity to the leaf’s middle vein. The arrival of the Ca^2+^ wave manifested as a maximum in the R-GECO1-mTurquoise ratio. Through determination of the distance between individual ROIs and the time difference between the ratio maxima, the speed of the waves was calculated.

### BiFC analysis

For transient expression in 5-6 week old *N. benthamiana* epidermal cells, *Agrobacterium tumefaciens* GV3101 (pMP90) carrying BiFC constructs were co-infiltrated with the p19 strain into leaves as described previously(*5*). The YFP C-terminal fragment SPYCE(M) was fused to the N-terminus of RBOHD and the N-terminal YFP fragment SPYNE(R)173 was fused to the N-termini of all 26 Arabidopsis CIPKs(*6*). Microscopic analyses of lower epidermal cells were conducted at 3 days after infiltration. An inverted fluorescence microscope, Leica DMI6000B equipped with a Leica N Plan L 20×/0.4 CORR PH1 objective (Leica, Wetzlar, Germany) and a Hamamatsu Orca camera (model C4742-80-12AG, Hamamatsu Photonics, Shizuoka, Japan) and a YFP filter set, which was operated with Openlab 5.0.2 software (Improvision, Coventry, UK) was used for BiFC quantification at lower magnification. For subcellular localization studies at higher magnification, an inverted confocal laser scanning microscope, Leica DMI6000, equipped with a Leica TCS SP5 II confocal laser scanning device (Leica Microsystems) and a 63×/1.2 water immersion objective (HCX PL APO lambda blue 63.0 × 1.20 Water UV) was used.

### Protein expression for *in vitro* assays

For protein purification, the sequences encoding the substrate proteins RBOHD N-terminus (aa 1-372), the inactive CPK5^D221A^ variant and the inactive CIPK26^K42N^ were fused with an N-terminal 2xStrepII-GST tag, the sequence of CBL1 was fused with an N-terminal 2xStrepII tag in the pET-24b vector (Merck, Darmstadt, Germany). Sequences of active CPK5 and CIPK26 were fused with an N-terminal 2xStrepII tag in the pIVEX-1.3WG vector (Biotechrabbit, Berlin, Germany). CPK5 and CIPK26 were expressed using the cell-free wheat germ RTS 500 system according to the manufacturer’s protocol (Biotechrabbit, Berlin, Germany). Substrate proteins were induced and expressed in *Escherichia coli* BL21 CodonPlus(DE3)-RIL cells (Stratagene) over night at 18°C after induction with 1 mM IPTG. Cell pellets were harvested, solubilized using a French press (Avestin EmulsiFlex-C3; ATA Scientific, Taren Point, Australia) and subsequently purified using Strep-Tactin-Macroprep (IBA-lifesciences, Germany). For the preparation of the 2xStrepII-GST-RBOHD-Nt protein, after mechanical cell disruption using the French-press, proteins were solubilized from inclusion bodies using 8 M urea (Drerup et al., 2014). StrepII-tagged proteins were purified using Strep-Tactin MacroPrep (IBA Lifesciences, Göttingen, Germany) following the manufacturer’s instructions.

Purified proteins (50 ng of CPK5 and 100 ng of CIPK26, 200 ng of CBL1, and 2000 ng of RBOHDs, 2000 ng of GST, 2000 ng of CIPK26K42N, 2000 ng of CPK5D221A) were mixed in reaction buffer (150 mM NaCl, 50 mM HEPES (pH 7.5), 0.5 mM Brij-35, 0.5 mM CaCl_2_, 5 mM MnS0_4_, 2 mM dithiothreitol, 10 µM Adenosine triphosphate (ATP), and 4 µCi of [γ-32P] ATP (3000 Ci mmol^- 1^). Reactions were incubated at 30°C for 30 min, stopped by addition of 6 µL 5 × SDS-loading buffer (125 mM Tris/HCl (pH 6.8), 5% (v/v) glycerin, 1% (w/v) SDS, 2.5% (v/v) β-mercaptoethanol, 0.025% (w/v) bromphenol blue) and analyzed by SDS-PAGE. Protein bands were fixed by Coomassie staining, and [γ-32P]-labeled proteins were visualized by autoradiography.

### ROS measurements in HEK293T cells

Vectors and methods for HEK293T cell transfection used in this study have been previously described(*7, 8*). The coding sequence of RBOHD was amplified by PCR and integrated into the pEF1-2xStrepII-N vector(*7*) and the pGGHEK vector(*8*). HEK293T cells were seeded into 96 well plates and incubated in (Fisher Scientific, Pittsburgh, USA) supplemented with 10% FBS (Fisher Scientific, Pittsburgh, USA) until reaching about 40% confluency. Transfection with plasmids carrying the coding sequences of the indicated proteins was performed using Genejuice transfection reagent (Merck, Darmstadt, Germany). 48 hours after transfection, ROS measurements were conducted as previously described(*7*). In brief, cells were subjected to a buffer containing horseradish peroxidase and L-012. Ionomycin was added to induce Ca^2+^ influx into the cells. ROS production was detected through luminescence measurements using either a Mithras2 LB943 (Berthold, Bad Wildbad, Germany) or a Tecan SPARK (Tecan, Männedorf, Switzerland) plate-reader.

### Ca^2+^ measurements in HEK293T cells using FURA-2

HEK293T cells were seeded into 96 well plates and incubated in D-MEM/Ham’s F-12 in D-MEM/Hams F-12 (FisherScientific, Pittsburgh, USA) supplemented with 10% FBS (Fisher Scientific,Pittsburgh, USA) until reaching about 80% confluency. Prior to the measurement, the media was aspirated from the cells and replaced with the Fura2-loading solution (5 µM Fura-2-AM and 0.15% Pluronic F127 (Invitrogen, Waltham, USA) dissolved in D-MEM/Ham’s F-12 with 10 %FBS). After one hour of incubation at 37°C at 5% CO_2_, the loading solution and cells were washed with HBSS -Ca^2+^, -Mg^2+^ (FisherScientific, Pittsburgh, USA). HBSS with indicated Ca^2+^ concentrations was added to the wells. Fura-2 fluorescence was measured with a Tecan Safire-2 plate reader (Tecan, Männerdorf, Switzerland). For the determination of absolute Ca^2+^ concentrations, Fura-2 fluorescence was calibrated with defined Ca^2+^ buffers generated with the Ca^2+^ Calibration Kit #1 (Invitrogen, Waltham, USA).

### Generation of HEK293T protein samples for LC-MS/MS analysis

HEK293T cells were cultivated in individual T-75 flasks (Sarstedt, Nümbrecht, Germany). At 30-50% confluency, cells were transfected with plasmids encoding the respective heterologous proteins using the Genejuice transfection reagent (Merck, Darmstadt, Germany). After 48 hours of incubation at 37°C at 5% CO_2_, media was aspirated, cells were washed with HBSS - Ca^2+^, -Mg^2+^ (FisherScientific, Pittsburgh, USA). After 5 minutes of incubation in HBSS + 125 µM Ca^2+^Cl_2_, 1 µM Ionomycin was added to induce Ca^2+^ influx. After 5 minutes, media was aspirated, ice cold PBS buffer was added, and cells were collected using a cell scraper. After one wash step in ice-cold PBS, cells were pelleted by centrifugation, the supernatant was aspirated, and cell pellets were flash-frozen in liquid nitrogen and stored at -80°C till further protein purification.

### LC-MS/MS-based quantitative proteome analyses

Proteins were extracted from cell pellets, and further sample processing and LC-MS/MS data acquisition were performed as described previously(*9*). Briefly, proteins were extracted and digested using a modified filter-assisted sample preparation protocol. After reduction and alkylation, proteins were digested using trypsine. For total proteome analysis, 10 μg of each sample were put aside and analysed without further processing. For phosphopeptide enrichment, 500 μg of peptides were enriched on titanium dioxide (TiO_2_)(*10*). LC-MS/MS analysis was performed by using an EASY-nLC 1200 (Thermo Fisher, Waltham, USA) coupled to a Q Exactive HF mass spectrometer (Thermo Fisher, Waltham, USA). Separation of peptides was performed on 20 cm frit-less silica emitters (New Objective, 0.75 µm inner diameter), packed in-house with reversed-phase ReproSil-Pur C_18_ AQ 1.9 µm resin (Dr. Maisch, Ammerbuch-Entringen, Germany). The column was constantly kept at 50 °C. Peptides were eluted in 115 min applying a segmented linear gradient of 0 % to 98 % solvent B (solvent A 0 % ACN, 0.1 % FA; solvent B 80 % ACN, 0.1 % FA) at a flow-rate of 300 nL/min. Mass spectra were acquired in data-dependent acquisition mode according to a TOP15 method. MS spectra were collected by the Orbitrap analyzer with a mass range of 300 to 1759 m/z at a resolution of 60,000 FWHM, maximum IT of 55 ms and a target value of 3×10^6^ ions. Precursors were selected with an isolation window of 1.3 m/z, and HCD fragmentation was performed at a normalized collision energy of 25. MS/MS spectra were acquired with a target value of 10^5^ ions at a resolution of 15,000 FWHM, maximum injection time of 55 ms and a fixed first mass of m/z 100. Peptides with a charge of +1, > 6, or with unassigned charge state were excluded from fragmentation for MS^2^, dynamic exclusion for 30 s prevented repeated selection of precursors.

Processing of raw data was performed using the MaxQuant software version 1.6.17.0 (Cox and Mann, 2008). MS/MS spectra were assigned to the uniport Homo Sapiens reference proteome supplemented with the sequences of the kinases used for transfection. During the search, sequences of 248 common contaminant proteins as well as decoy sequences were automatically added. Trypsin specificity was required and a maximum of two missed cleavages was allowed. Carbamidomethylation of cysteine residues was set as fixed, oxidation of methionine, deamidation and protein N-terminal acetylation was set as variable modifications. A false discovery rate of 1% for peptide spectrum matches and proteins was applied. Match between runs and requantify options were enabled.

